# Bayesian AMMI-Based Simulation of Genotype × Environment Interactions

**DOI:** 10.64898/2026.03.11.711188

**Authors:** Heegun Lee, Vitor Seiti Sagae, Julian Garcia-Abadillo, Fernando Bussiman, Mitzilin Zuleica Trujano-Chavez, Jorge Hidalgo, Diego Jarquin

## Abstract

Genotype-by-environment interaction (GEI) has been studied to identify environment-stable/favorable genotypes. The GEI simulation could help refine the inference by incorporating tangible factors such as genomic and environmental information. The Bayesian additive main effect and multiplicative interaction (Bayesian AMMI) model captures the genotype-specific responses across environments, reflecting directional relationships between genotypes and environments. Thus, we propose a Bayesian AMMI-based GEI simulation framework that utilizes high-throughput environmental covariance matrices to generate GEI effects with interpretable directional structure. To demonstrate the proposed approach, two simulated phenotypes were assessed under four levels of GEI variance. In the first simulation (Sim1), GEI effects were sampled from a multivariate normal distribution defined by the GEI matrix. In the second simulation (Sim2), GEI effects were generated by extending Sim1 with the Bayesian AMMI model. In both simulations, increasing GEI variance resulted in lower correlations of phenotypes across environments and stronger genotype-specific sensitivity to environmental variation. Across five cross-validation designs, models accounting for GEI consistently outperformed one that did not, with prediction accuracy generally decreasing as GEI variance increased. Clear distinctions between the two simulated phenotypes were evident from biplot analyses: Sim2 successfully captured environmental relatedness and genotype-specific responses, whereas such structure was absent in Sim1. These results demonstrate that the proposed Bayesian AMMI-based GEI simulation framework enables interpretable visualization of GEI and supports genomic selection strategies under complex environmental conditions.

## Introduction

The genotype-by-environment interaction (GEI) has been studied in animal and plant breeding aiming to select genotypes that perform consistently (stably) across environments or favorably (locally adapted) in specific environments. Visualization of GEI patterns using biplots is widely applied in plant breeding as an approach to identify genotypes relevant to environments (Fonseca et al. 2022; Jarquín et al. 2016). Recently, Kebede et al. (2024) applied this approach in chickens to explore breed-by-environment interactions, aiming to optimize breeding strategies for smallholder farms. This work highlights the potential for transferring GEI visualization methodologies developed in plant breeding to animal breeding contexts. However, evaluation of genotype performance across diverse and potentially unobserved environments remains challenging. In this sense, simulations for evaluation of genotype performance across environments, accounting for interpretable visualization information have become crucial. Existing studies utilizing GEI simulations frequently adopt simplified environmental models, which potentially limits their applicability to complex, multi-faceted ecosystems (Liu et al. 2024; Majumdar et al. 2021; Verhulst 2025). However, the actual GEI is likely a genotype’s response to complex environmental conditions. To account for complex environmental conditions in GEI simulation, we propose the use of high-dimensional environmental covariates within a Bayesian additive main effect and multiplicative interaction (Bayesian AMMI) model.

The AMMI model is a method widely used in plant breeding to characterize GEI patterns between genotypes and environments (Gauch 1988). This combines analysis of variance for additive effects and singular value decomposition (SVD) for non-additive effects (Gauch 1992). This approach enables the visualization of GEI in biplot which provides directional relationships among genotypes and environments. These directional relationships illustrate the extent of genotype stability or their adaptability to specific environments, thereby assisting breeders to intuitively select desirable genotypes. However, SVD in AMMI accounting for GEI may lead to biased interpretations when data contain outliers (Rodrigues et al. 2015).

To address this constraint, Bayesian treatment of AMMI was introduced to allow inference-based estimation of interaction parameters (Crossa et al. 2011; Viele and Srinivasan 2000). Especially as a solution to overparameterization in AMMI, Crossa et al. (2011) proposed using the von Mises-Fisher (VMF) distribution in the Gibbs sampler to simulate the interaction parameters. This approach allows for the derivation of credible intervals for the GEI parameter based on directional relationships between genotypes and environments. In this sense, employing appropriately simulated genotypic and environmental data as prior information, using the VMF distribution, holds the potential to generate new GEI values that incorporate directional information of GEI.

Simulation of environmental variables can be achieved through environmental covariates. Jarquín et al. (2014) introduced covariance structures to account for main and interaction effects of markers and environmental covariates, addressing computational challenges associated with high-dimensional environmental data. In the same way as the genomic relationship matrix (GRM), an environmental covariance matrix can be constructed using environmental covariates, and environmental effects can be represented as random variables drawn from this matrix. Extending this concept, incorporating the environmental covariance matrix into GEI modeling allows for simulation of GEI variables that reflect environmental conditions.

Therefore, the objective of this study was to propose a simulation framework for GEI that employs the Bayesian AMMI model utilizing an environmental covariance matrix. The framework enables GEI simulation for environments characterized by complex, high-throughput environmental covariates. We evaluated two simulation scenarios, each with 50 replicates, under varying GEI variance levels to assess the effectiveness of the proposed method across different interaction magnitudes. To demonstrate the utility of this framework, we analyzed patterns in phenotypic performance across environments, SNP effect estimates, genomic prediction accuracy, and graphical GEI interpretation through biplots with AMMI stability values (ASVs).

## Materials & Methods

The simulation was performed in two steps: 1) pre-simulate phenotypes, and 2) rescale GEI effects based on pre-simulated phenotypes. The simulation workflow is provided in Figure 1. Generation of founder population was performed using the ‘runMacs’ command in AlphaSimR (Gaynor et al. 2020), and recent population was generated using in-house developed R pipeline based on gene drop approach. Through these simulation steps, a genotype file containing allele dosages (0, 1, or 2) along with pedigree and phenotype files were generated. The complete pipelines for simulation, the corresponding command lines, and an example of parameter settings are provided in Supplementary Files 1, 2, and 3, respectively.

**Figure 1.**
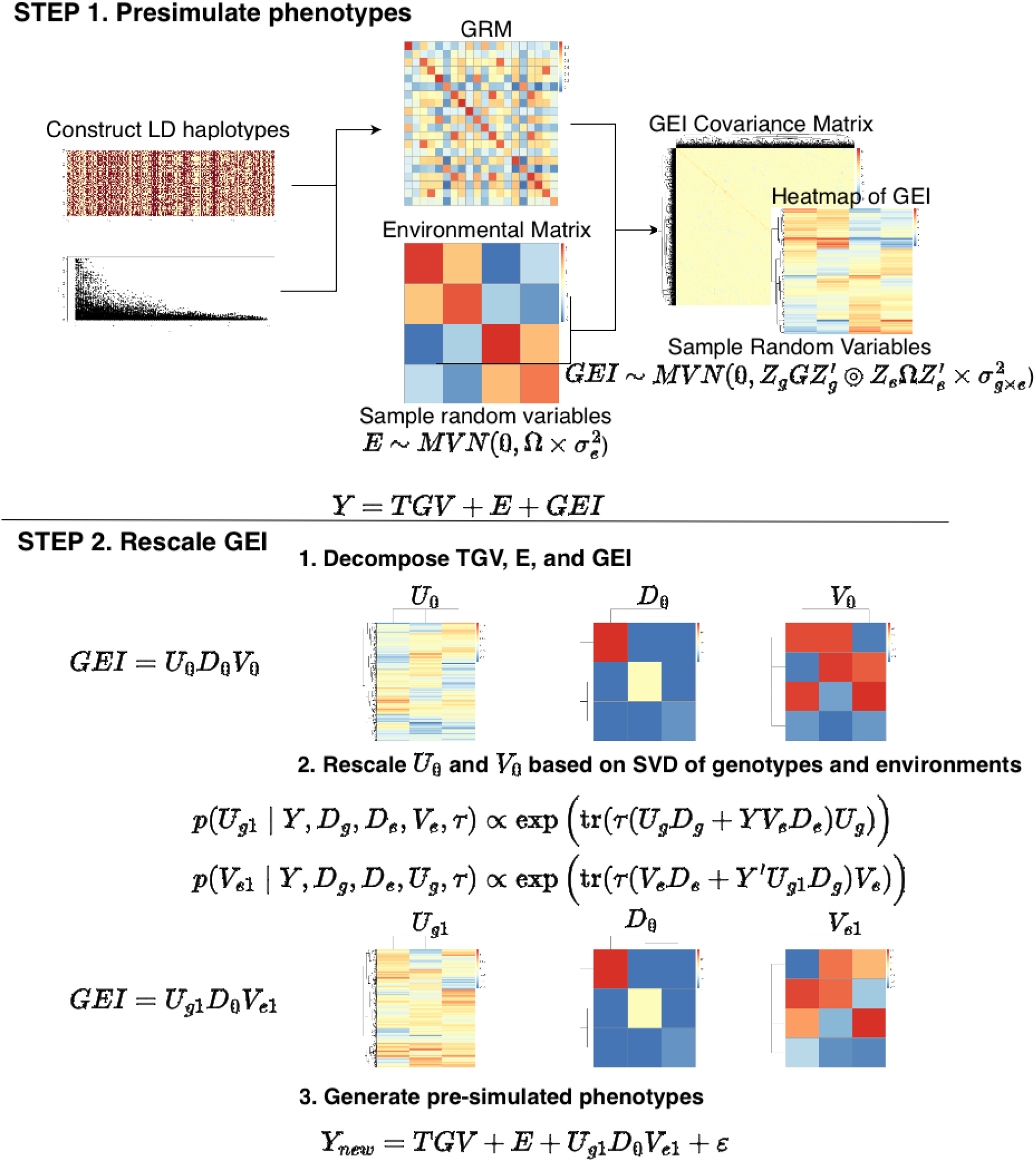
Overview of framework of GEI simulation applying Bayesian AMMI rescaling method. The diagram illustrates the main steps of the simulation framework, including the construction of genomic and environmental covariance structures, simulation of genotypes and environments, and generation of phenotypic values. STEP1 presents the process of pre-simulating variables. STEP2 presents rescaling steps and change of trends in singular vector matrices after rescaling.

In the first step, the linkage disequilibrium (LD) haplotypes used as the founder population were generated under the following conditions using AlphaSimR: a cattle demographic history (species = ‘CATTLE’), a total of 500 individuals (nInd = 500), and 2,500 markers (segSites = 2,500) on one chromosome (nChr = 1). Simulations were conducted using a single chromosome to reduce computational burden (Honorato-Mauer et al. 2024).

Afterwards, a subset of the generated alleles was classified as *m* causal alleles directly underlying true quantitative trait loci (QTL) effects, and the remaining alleles were treated as alleles in LD with these QTL. The initial *m* causal allele effects were generated as **qtl**_**0**_ = {*qtl*_*m*_} ~ Gamma(*t, θ*) where the shape parameter *t* was defined by the user and *θ* was fixed at 1. Causal allele effects were then rescaled to match genetic variance to be the user-defined value, using 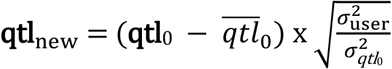, where 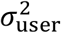 is the target (user-defined) genetic variance at the QTL level, and 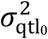 is the variance of the initially simulated causal allele effects before scaling (Sargolzaei and Schenkel 2009).

### Simulate recent population (genotype)

Afterwards, those alleles were inherited across recent generations in the population using a gene drop approach, considering crossover and mutation events at each meiosis, thereby capturing LD changes over time (MacCluer et al. 1986). Crossover events were modeled using a Poisson distribution, 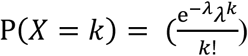, where *X* is the number of crossovers in a given interval, *k* is a specific possible number of crossover events, and *λ* is the product of the distance between loci (in centimorgans, cM) and the cross over probability (Sargolzaei and Schenkel 2009). The distance between loci was calculated as the chromosome length divided by the number of loci.

### Simulate recent population (phenotype)

Prior phenotypes were then generated as the sum of genomic, environmental, and GEI effects, and subsequently used as input for the Bayesian AMMI Gibbs sampler algorithm during the rescaling process. The phenotype of the *i*-th genotype (*i* = 1, 2, …, *n*_*g*_) in the *j*-th environment (*j* = 1, 2, …, *n*_*env*_) was expressed as:

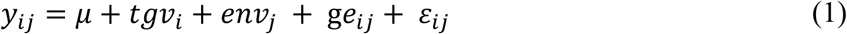

where *y*_*ij*_ is the phenotype of *i*-th genotype in *j*-th environment; µ is a common effect; *tgv*_*i*_ represents the true genetic value of *i*-th genotype, determined by the effects of inherited causal alleles from the previous generation; *env*_*j*_ is a random variable sampled from a multivariate normal distribution defined by the environmental covariance matrix 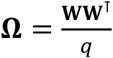, where **W** is an *n*_*env*_ × *q* matrix of user-defined environmental covariates, and *q* is the number of covariates; thus, 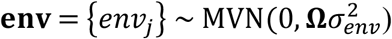, the GEI term, ge_ij_ represents the interaction of the *i*-th genotype and the *j*-th environment, sampled as 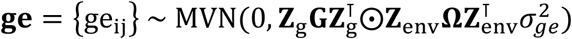, where **G** is the GRM, **Z**_g_ and **Z**_env_ represent the incidence matrices for genotypes and environments, respectively, and ⨀ denotes the Hadamard product. The random residuals *ε*_*ij*_ were independently sampled as 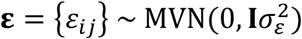. The variances for **tgv, env, ge**, and **ε** are defined by the user. In this study, four levels of GEI variance (0.1, 0.5, 1.0, and 2.0) were evaluated, while the genetic, environmental, and residual variances were fixed at 0.5, 0.5, and 0.1, respectively.

Through this step, **env** and **ge**, which depend on environmental covariates, were generated. However, **ge** at this stage reflects the interaction between genotypes and the general environmental relationship, rather than accounting for the specific environmental conditions in the given generation, since it follows the assumption 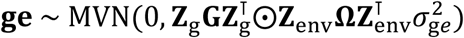.

### Rescale GEI variables using Gibbs sampler algorithm

As a subsequent step, **ge** variables were updated to incorporate the effects of the given generation’s **tgv** and **env** components. This update was performed using the Gibbs sampler algorithm as implemented in the Bayesian AMMI.

The Bayesian AMMI, as described by Crossa et al. (2011), incorporates inferential statistics and a standard set of priors into the AMMI model to ensure that model constraints and conditional conjugacy are satisfied. The joint posterior inference for singular vectors of GEI matrix for genotypes and environments, which follows the VMF distribution, can be obtained by iteratively sampling from the following conditional distributions (Hoff 2009):

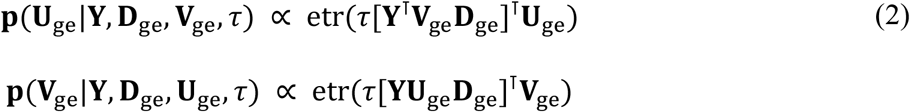

where **Y** is the *n* × *p* phenotype matrix representing *n* samples from a *p*-variate (environments) distribution; **U**_ge_, **V**_ge_, and **D**_ge_ denote the orthogonal matrices of SVD corresponding to the genotypic, environmental, and singular value components of the GEI effects, respectively; *τ* represents the precision, defined as the reciprocal of the residual variance, 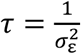. The operator etr denotes the exponential of the trace.

To extend the Bayesian AMMI to the GEI simulation, Equation (1) can be expressed in matrix notation as:

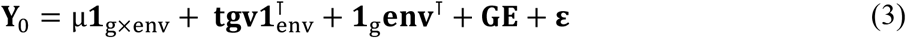

where µ is a common effect and **1**_g×env_ is a matrix of ones of order 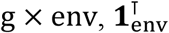 and **1**_g_ are vectors of ones used to expand **tgv** and **env**^⊺^ to matrices. The subscript ‘0’ on **Y** denotes the pre-simulated phenotypic performance. After applying SVD to 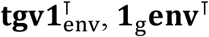, and **GE**, the formula can be represented as:

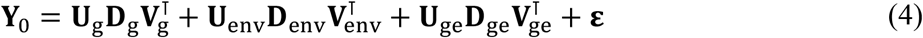

where **U**_∗_ and **V**_∗_ are orthogonal matrices of left and right singular vectors, respectively; **D**_∗_ is a diagonal matrix of singular values; and subscripts g, env, and ge indicate the genomic, environmental, and GEI components, respectively.

Based on these derivations, we formulated a new expression to model GEI that captures directional relationships by incorporating both 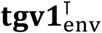 and **1**_g_ **env**^⊺^. This assumed that the effects of genotypes (or environments) can be decomposed into GEI-relevant and GEI-irrelevant components. The conditional posterior distributions for updating the corresponding genotype and environment singular vectors associated with GEI are then defined as follows:

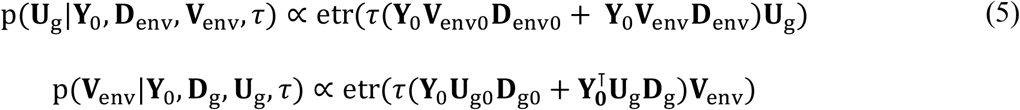

where **Y**_0_**V**_env0_**D**_env0_ (or **Y**_0_**V**_g0_**D**_g0_) represents the portion of the phenotypic variation that is not attributed to environmental (or genotypic) effects, whereas 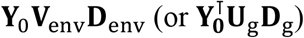 represents the portion of phenotypic variation that is attributed to environmental (or genotypic) effects. Since ***tgv*** and ***env*** were independently generated in the pre-simulation steps, **Y**_0_**V**_env0_**D**_env0_ and **Y**_0_**U**_g0_**D**_g0_ can be redefined as **U**_g_**D**_g_ and **V**_env_**D**_env_, respectively. This allows Equation (5) to be written as:

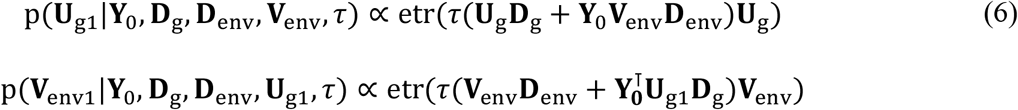

Combining all these assumptions, (3) can be updated to

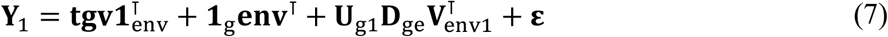

where **U**_g1_ and **V**^⊺^ are rescaled singular vectors via Bayesian AMMI Gibbs sampler algorithm using pre-simulated data as prior information as the process described above, and the **Y**_1_ represents the phenotype matrix after Bayesian AMMI-based rescaling of GE effects.

### Parameters for recent population used in this study

Table 1 provides an overview of parameter settings used for biological events during meiosis, the mating design, and the recent population simulation. Four distinct environments, denoted as E1, E2, E3, and E4 were simulated in this study. To clearly illustrate GEI trends, monthly records of temperature at 2 meters (T2M) for these environments were obtained from NASA POWER database for the year 2023 (Sparks 2018). Based on the 2023 T2M, the four environments were classified into two distinct climate types: E1 and E2 were positively correlated, as were E3 and E4, while the two climate types were negatively correlated with each other. The environmental covariance matrix used in this study was defined as:

**Table 1.**
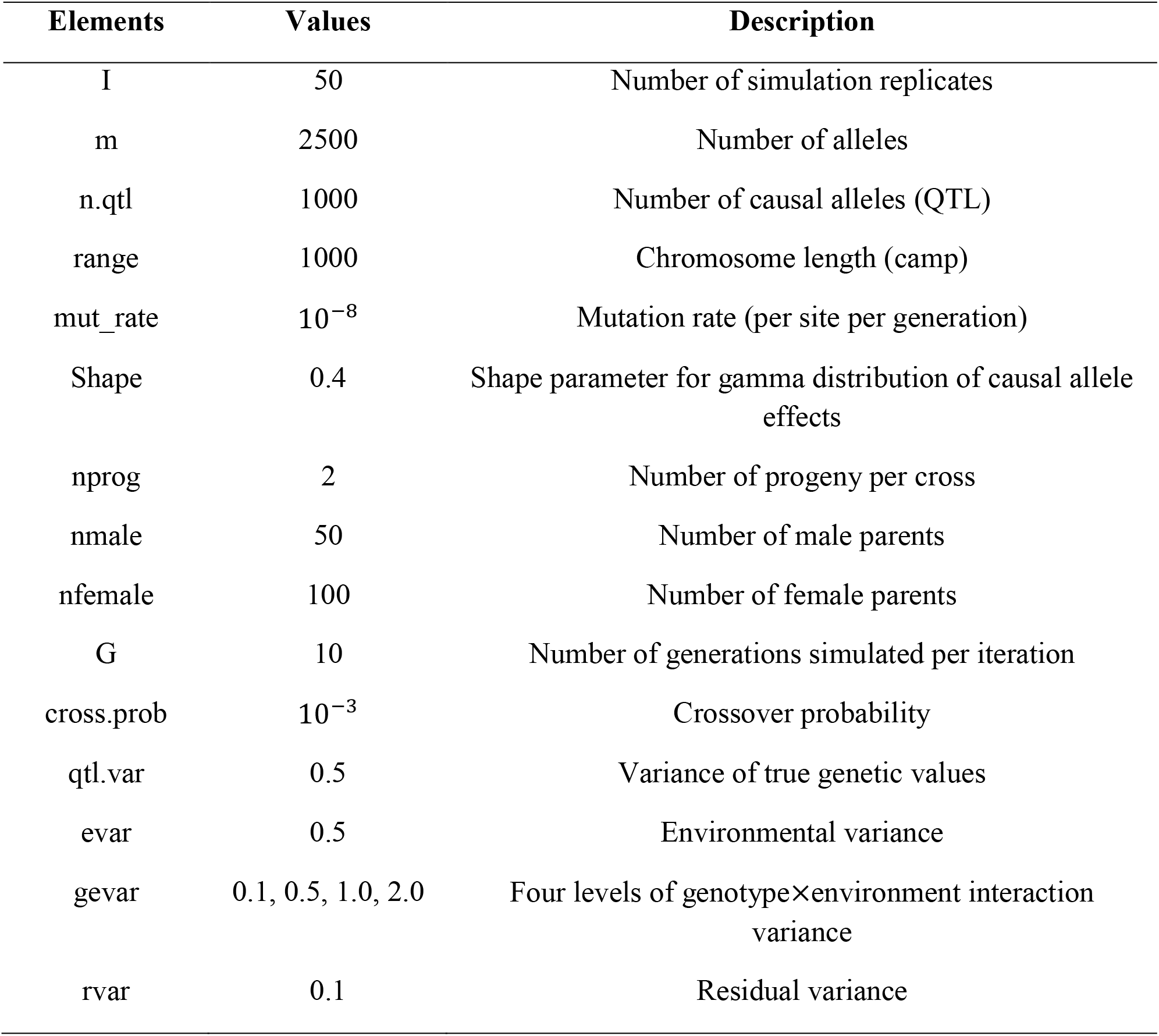
Parameters used for simulation of the recent population in this study. To assess the effect of genotype × environment interaction (GEI) on phenotypic variation, four levels of GEI variance (0.1, 0.5, 1.0, 2.0) were applied across simulation replicates. Other simulation parameters, including the number of markers, number of causal alleles, chromosome length, population structure, mutation rate, and environmental/residual variance, were set as indicated.

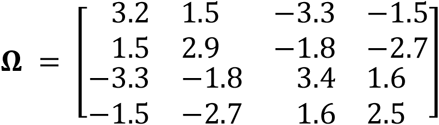

The monthly T2M records utilized in this study are represented in Table 2. Users may incorporate different types of environmental covariates depending on their research objectives. Based on these parameters, 50 replicates were conducted using randomly selected individuals over 10 generations, each involving the mating of 50 males with 100 females, resulting in two progenies per female.

**Table 2.** Monthly records of temperature at 2 meters (T2M, ℃) from four locations in 2023 used in this study. Data were obtained from NASAPOWER database. E1 and E2 shared similar environmental conditions, whereas E3 and E4 exhibited a different but internally consistent environmental trend.

| T2M monthly records |  |  |  |  |  |  |  |  |  |  |  |  |
| --- | --- | --- | --- | --- | --- | --- | --- | --- | --- | --- | --- | --- |
|  | Jan | Feb | Mar | Apr | May | Jun | Jul | Aug | Sep | Oct | Nov | Dec |
| E1 | -1.78 | 1.94 | 4.66 | 13.18 | 13.18 | 16.28 | 22.17 | 25.61 | 24.16 | 15.82 | 8.42 | -0.42 |
| E2 | 0.68 | -0.17 | 1.49 | 7.49 | 11.12 | 15.88 | 20.58 | 22.96 | 20.14 | 14.76 | 7.33 | -0.27 |
| E3 | 25.56 | 25.03 | 22.64 | 19.53 | 15.92 | 12.39 | 11.55 | 16.53 | 18.46 | 21.24 | 23.70 | 23.51 |
| E4 | 23.85 | 23.87 | 23.46 | 22.63 | 20.90 | 18.84 | 17.26 | 18.47 | 22.59 | 21.63 | 22.05 | 22.80 |

### Demonstration of simulation

To demonstrate how effectively the Bayesian AMMI framework can be applied for GEI simulation, two types of simulation implemented through Equations (3) and (7) were evaluated. In the first simulation (Sim1), GEI effects were sampled from a multivariate normal distribution defined by the GEI matrix. In the second simulation (Sim2), GEI effects were generated by extending Sim1 with the Bayesian AMMI framework (Equation 7).

### Phenotype trends across the environments

Simulated GEI trends across the environments were assessed by examining phenotypic pairwise correlations. Additionally, trends in SNP solutions across environments were evaluated to determine whether the predicted SNP effects captured GEI patterns appropriately. The SNP solutions for each environment were obtained by back solving the genomic estimated breeding values (GEBVs) estimated separately for each environment using postGSf90 (Lourenco et al. 2022). The single-trait mixed linear model used to obtain GEBVs was specified as:

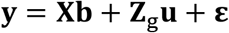

where **y** is the vector of phenotypic records; **b** is the vector of fixed effects (sex and generations); **u** is the vector of additive genetic random effects with 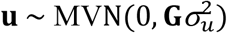, where 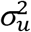 is the additive genetic variance; and **X** and **Z**_g_ are the incidence matrices for fixed and random effects, respectively.

The phenotypic trends were further assessed by examining changes in regression coefficients within the classification. Genotypes were approximately classified into two groups based on their response patterns: an ascending group and a descending group. A genotype was categorized as ascending group if it exhibited the lowest performance in E1 or E2, and as descending group if the lowest performance was observed in E3 or E4. The trends of regression coefficients across environments were then analyzed for each GEI variance. To quantify the impact of environments on phenotypes, regression coefficients were estimated using the model:

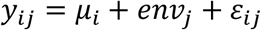

where *μ*_*i*_ is the mean performance of the *i*th genotype across environments. To estimate the regression coefficients, the environments were temporarily coded as original values from 1 to 4. This classification assumed that individuals exhibiting similar GEI trends are likely to possess comparable adaptability to environments with similar climate conditions. In other words, each genotype’s performance could differ in its interaction with environments, reflecting variation in GEI.

### Prediction accuracy

Three prediction models were compared to evaluate the impact of GEI variance on prediction accuracy. The first model (M1) included only the main effects and was specified as:

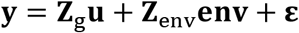

Two additional models (M2 and M3) explicitly incorporated **GEI** effects:

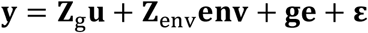

Both M2 and M3 models were fitted using a Bayesian mixed model implemented in the BGLR R package (Pérez and de los Campos 2014). A total of 20,000 iterations were used, with a burn-in period of 2,000. The covariance structures assumed for the GEI effects followed Equations (3) and (7), respectively.

Prediction accuracy was assessed as the Pearson’s correlation between phenotype and the corresponding sum of the predicted solutions for those effects. A single replicate of a five-fold cross-validation 1 (CV1; Predicting performance of new developed lines through relationships with others) was implemented for each iteration set.

### AMMI stability value

The stability trends of genotypes were also compared between the two different phenotypes, Sim 1 and Sim2. The ASVs, representing the scaled distance from the origin to each genotype’s coordinates in the biplot, was used as a stability indicator. In practice, ASV is used to quantify the stability of genotypes across the environments, and its reliability depends on the accuracy of the biplot on which it is based (Pour-Aboughadareh et al. 2022; Zali et al. 2012). It accounts for the magnitude of each principal component axis’s contribution to the GEI, providing a quantitative measure of genotype stability (Purchase et al. 2000). The ASV is calculated as follows:

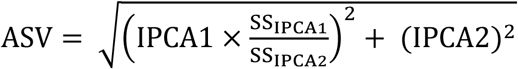

where IPCA1 and IPCA2 are the first and second interaction principal component axes in the AMMI model, and SS_IPCA1_ and SS_IPCA2_ are the corresponding sums of squares, respectively. The correlation of ASV between Sim1 and Sim2 phenotypes was calculated to quantify the degree of similarity in the stability they represented.

## Results & Discussion

### Phenotypes and marker effects

The phenotypic correlations across the four simulated environments were compared to evaluate whether the simulated GEI effects reflected the environmental variability. Correlations among phenotypes generally reflected the imposed environmental trends, and their magnitude varied according to the specified GEI variance (Table 3). In both Sim1 and Sim2, phenotypic performance was increasingly influenced by environments as the GEI variance increased. Phenotypic correlations between similar environmental conditions (E1 and E2; E3 and E4) changed little across different GEI variance levels (Figure 2).

**Table 3.** Phenotypic correlation between pairs of environments under two simulation approaches (Sim1 and Sim2) and different levels of GEI. Values represent mean ± SD across 50 replicates. GEI variance indicates the simulated magnitude of GEI effects (0.1, 0.5, 1.0, 2.0).

| Envs<sup>1</sup> / GEI var | Sim1 |  |  |  | Sim2 |  |  |  |
| --- | --- | --- | --- | --- | --- | --- | --- | --- |
|  | 0.1 | 0.5 | 1.0 | 2.0 | 0.1 | 0.5 | 1.0 | 2.0 |
| E1-E2 | 0.89±0.04 | 0.82±0.05 | 0.75±0.06 | 0.68±0.05 | 0.83±0.05 | 0.74±0.07 | 0.67±0.09 | 0.58±0.11 |
| E1-E4 | 0.64±0.12 | 0.41±0.11 | 0.27±0.10 | 0.07±0.08 | 0.60±0.11 | 0.37±0.11 | 0.24±0.11 | 0.02±0.11 |
| E1-E3 | 0.50±0.16 | 0.16±0.10 | -0.05±0.10 | -0.31±0.08 | 0.48±0.13 | 0.21±0.11 | 0.03±0.11 | -0.18±0.08 |
| E2-E4 | 0.54±0.16 | 0.27±0.11 | 0.06±0.11 | -0.21±0.08 | 0.52±0.15 | 0.32±0.11 | 0.15±0.12 | -0.08±0.10 |
| E2-E3 | 0.60±0.14 | 0.36±0.11 | 0.18±0.12 | 0.01±0.08 | 0.56±0.13 | 0.34±0.12 | 0.18±0.13 | -0.02±0.10 |
| E3-E4 | 0.89±0.05 | 0.82±0.05 | 0.78±0.04 | 0.70±0.04 | 0.83±0.06 | 0.73±0.08 | 0.67±0.07 | 0.58±0.09 |

<sup>1</sup>Pairs of assessed environments (e.g., E1-E2, E1-E3, etc.)

**Figure 2.**
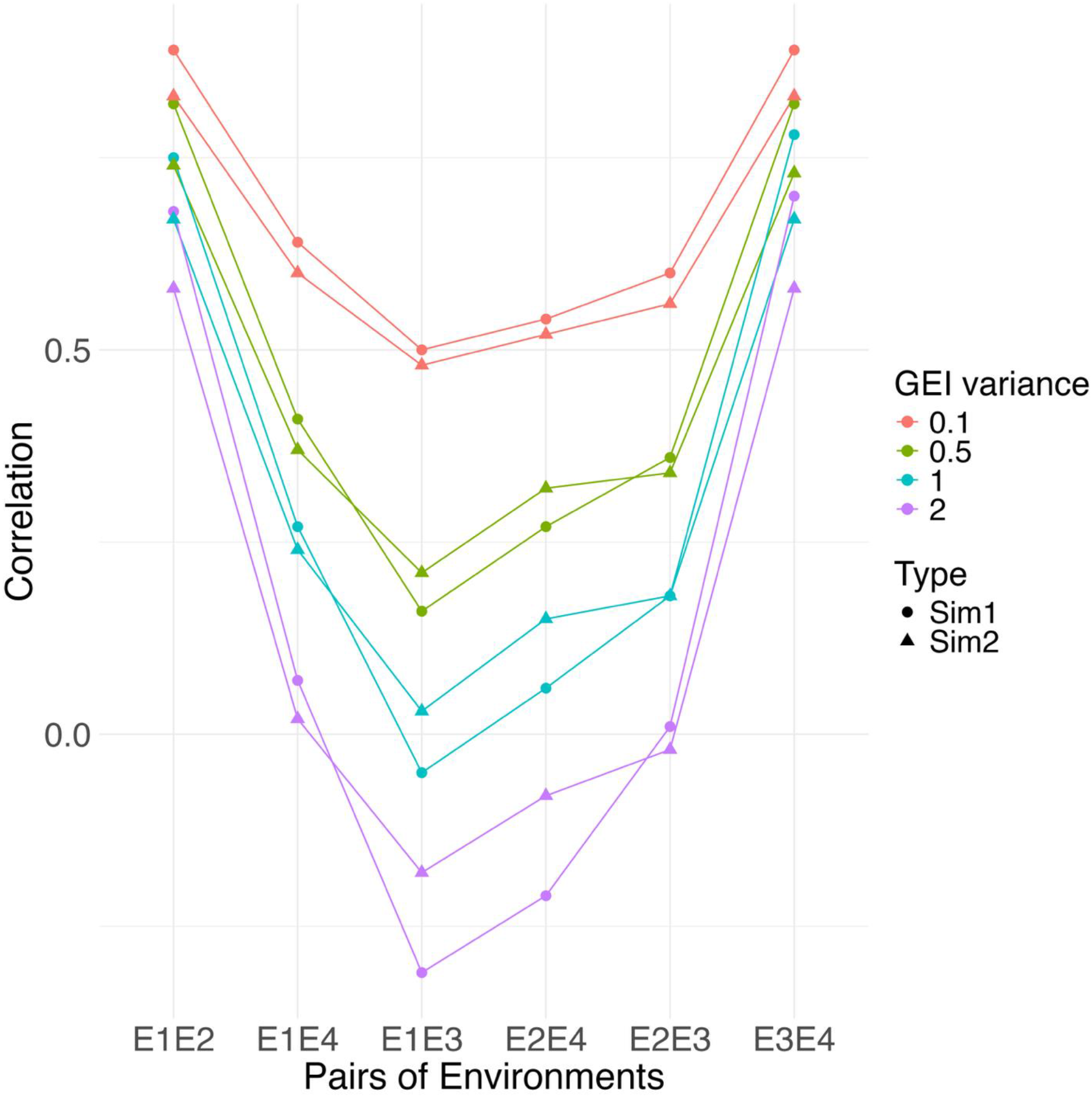
The correlation of phenotypic performances between the environments. Sim1 represents phenotypes including GEI generated following multivariate normal distribution, while Sim2 represents phenotypes including GEI rescaled through Bayesian AMMI approach.

Conversely, correlations between contrasting climate types fluctuated, which were close to or below 0 when the GEI variance was set to 1 or 2. As the GEI variance increased, the gap between correlations from similar climate trends (E1 and E2; E3 and E4) and those from contrasting climates (e.g. E1 to E3 or E4) widened. This trend is consistent with the study by Kolmodin and Bijma (2004), which stated that the higher selection response to environments was identified with a higher degree of GEI effects. Both Sim1 and Sim2 exhibited comparable overall phenotypic correlation patterns.

The trends of SNP solutions for correlations across environments were similar to those observed for phenotypes (results not provided). This was consistent with the role of GEI in determining the strength of relationship between genomic trends and environments. Similar with the phenotypic correlation comparisons, the numerical differences between Sim1 and Sim2 were not significant.

### Regression coefficient

In both phenotypes, the intensity of regression coefficients increased with GEI variance, accompanied by large standard deviations (SD;Table 4). These large SD likely reflected the genotype-specific nature of GEI in both phenotypes. Since the genotypes in this study were randomly selected, the expected response of genotypes to different environments could vary due to difference in the genomic structures of individuals within a population. The trend of variability in this comparison has also been observed in empirical data. In haplotype-by-environment studies, the environmental response of genotypes was evaluated at the haplotype levels (Chen et al. 2007; Patiranage et al. 2021). Fine-scale GEI studies in both animal and plant breeding have demonstrated the environmental responsiveness of individual genetic variants as well. For example, Smith et al. (2022) identified sixteen QTL regions associated with growth traits in U.S. Red Angus cattle through a genome-wide association study that accounted for GEI. Similarly, Li et al. (2018) reported significant QTL-by-environment interaction using multi-environment and multi-traits QTL analyses for drought tolerance in maize.

**Table 4.** The regression coefficients of phenotypes across different GEI variance parameters (0.1, 0.5, 1.0, 2.0) under two simulation approaches (Sim1 and Sim2). Values represent mean ± SD across 50 replicates.

| GEI var \ Group | Sim1 |  | Sim2 |  |
| --- | --- | --- | --- | --- |
|  | Ascending<sup>1</sup> | Descending<sup>2</sup> | Ascending<sup>1</sup> | Descending<sup>2</sup> |
| 0.1 | -0.17±0.03 | 0.17±0.04 | -0.30±0.10 | 0.31±0.09 |
| 0.5 | -0.54±0.11 | 0.53±0.10 | -0.69±0.17 | 0.67±0.17 |
| 1.0 | -0.85±0.21 | 0.82±0.20 | -1.01±0.32 | 0.99±0.28 |
| 2.0 | -1.38±0.26 | 1.34±0.27 | -1.19±0.40 | 1.16±0.34 |

<sup>1</sup>Ascending: individuals with the lowest performance in environments 1 or 2.
<sup>2</sup>Descending: individuals with the lowest performance in environments 3 or 4.

### Genomic prediction accuracy

The genomic prediction accuracy under different levels of GEI variance was evaluated across scenarios. The prediction accuracy from all models decreased as GEI variance increased (Table 5). Both M2 and M3 models exhibited better accuracy than M1, which didn’t account for GEI in the model. The M2 predicted phenotypes more accurately compared to M3 across all scenarios (Figure 3).

**Table 5.** The Pearson’s correlations between observed phenotypes and predicted values from three models. Values represent mean ± SE across 50 replicates.

| GEI var \ Models | M1<sup>1</sup> | M2<sup>2</sup> | M3<sup>3</sup> |
| --- | --- | --- | --- |
| 0.1 | 0.79±0.006 | 0.83±0.005 | 0.82±0.005 |
| 0.5 | 0.70±0.005 | 0.77±0.004 | 0.75±0.004 |
| 1 | 0.63±0.005 | 0.74±0.004 | 0.71±0.003 |
| 2 | 0.52±0.005 | 0.70±0.004 | 0.64±0.003 |

<sup>1</sup>M1: model including true genetic value (TGV) and environmental main effects (E).
<sup>2</sup>M2: model including TGV, E, and GEI effects simulated using the Sim1 phenotypes using a multivariate normal distribution.
<sup>3</sup>M3: model including TGV, E, and GEI effects simulated using the Sim2 phenotypes rescaled using the Bayesian AMMI framework.

**Figure 3.**
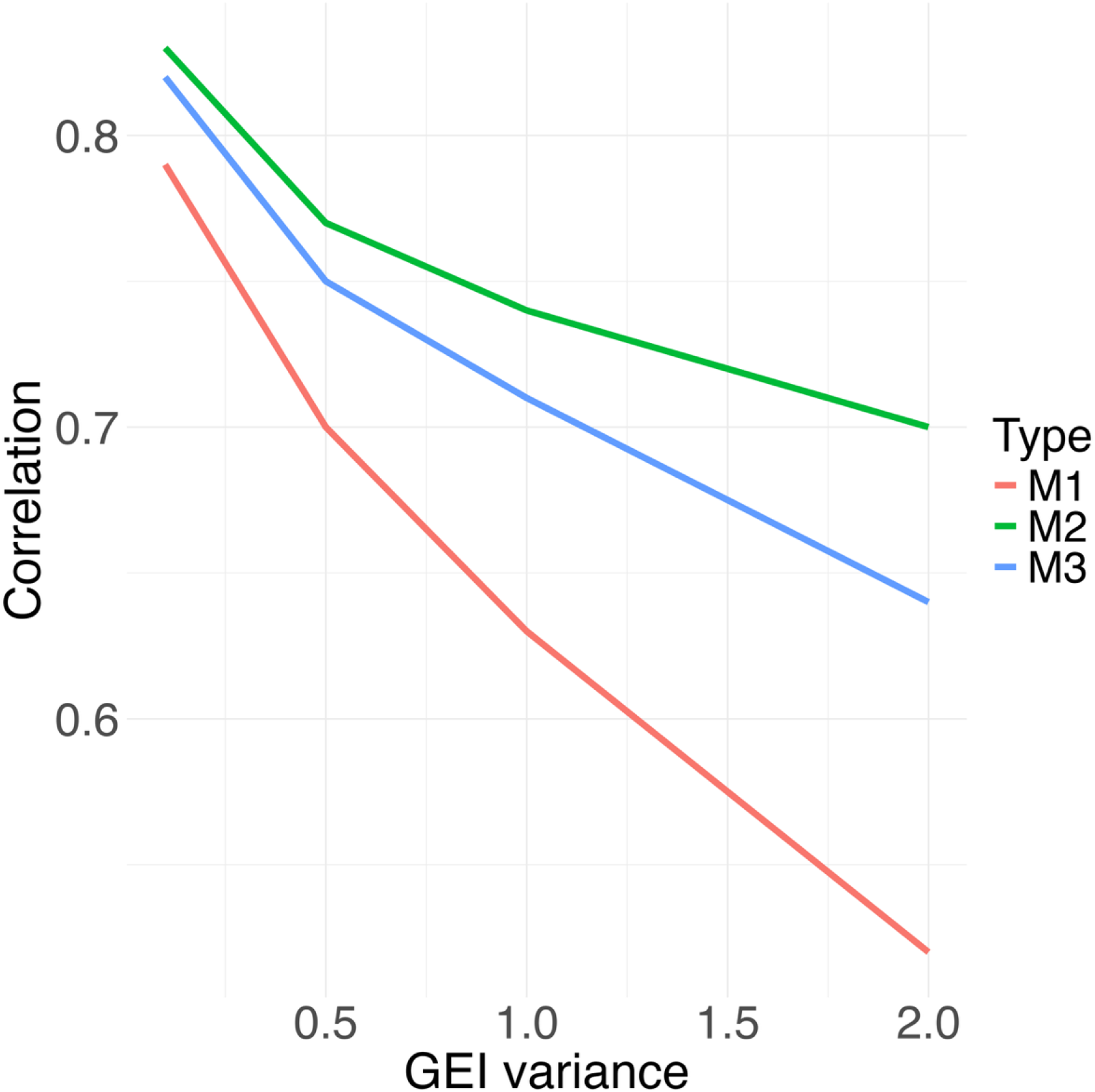
Comparison of prediction accuracy among models. M1: model including GRM and Ω; M2: model including GRM, Ω, and GE trained on Sim1 phenotypes, where GEI was generated using a multivariate normal distribution; M3: model including GRM, Ω, and GE trained on Sim2 phenotypes, where GEI was rescaled using the Bayesian AMMI framework.

The gap in accuracy between these models widened as GEI variance increased. The lower accuracy of M3 compared to M2 not necessary imply that M2 is an appropriate simulation approach. For both models M2 and M3, the GEI matrix was represented using the Hadamard product of the GRM and environmental covariance matrix. In M2, the GEI variables used to train the model were drawn from a multivariate normal distribution based on the GEI matrix. In contrast, the GEI variables in M3 did not exactly correspond to this Hadamard product which led to systematically lower prediction accuracy. Thus, differences in accuracy should not be interpreted as evidence that M2 biologically captures more GEI variance but instead arise from differences in how similarly the underlying variable matrices were simulated.

Overall, models incorporating GEI effects showed higher prediction accuracies compared to the one without considering GEI, consistent with Jarquín et al. (2014) which reported improved prediction accuracy for grain yield when GEI was included in the model.

### Biplot and directional relationship

Both Sim1 and Sim2 showed that the phenotypic trends were influenced by the simulated GEI variables in proportion to the magnitude of the GEI variance. Visual comparisons highlight the advantage of the Bayesian AMMI approach. Ten individuals from the first generation of one replicate were selected to visualize GEI in a biplot (Figure 4). In these results, the distances among the environments in the Sim1 did not reflect the directional relationships between environmental variables, whereas Sim2 did.

**Figure 4.**
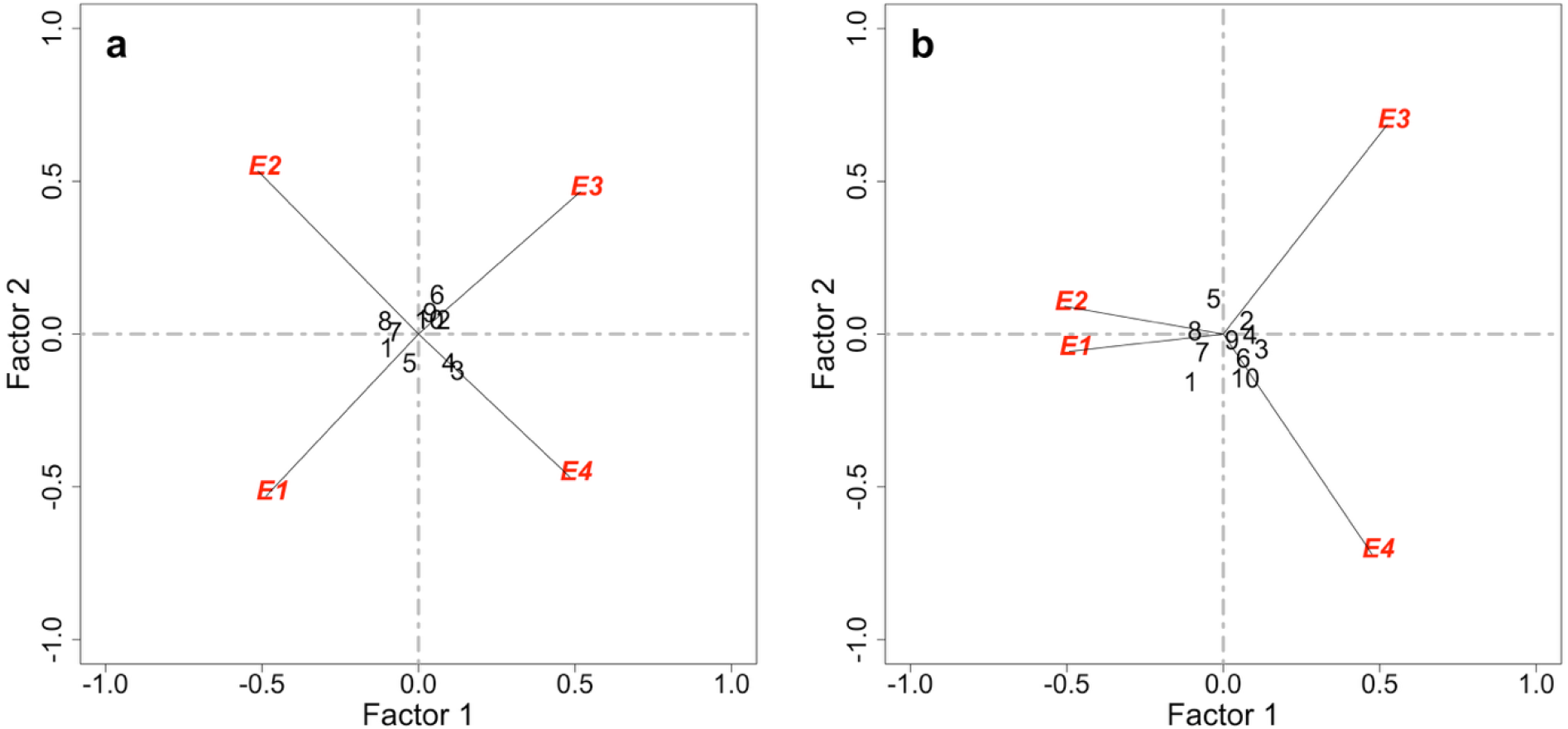
The biplot representing directional relationship of 10 randomly selected genotypes to four environments from generation 0. (a) The biplot using phenotypes including GEI generated following multivariate normal distribution (Sim1). (b) The biplot using phenotypes including GEI rescaled through Bayesian AMMI approach (Sim2).

For example, in one replicate from this study, the simulated environmental values were 0.38, 0.88, −0.54, and −0.73, for E1, E2, E3, and E4, respectively. Based on these values, E1 and E2, and E3 and E4 were supposed to be positioned closer together in the biplot, reflecting GEI relationships. However, for the Sim1, the environments were equally spaced apart from each other in the biplot, regardless of the simulated values. This is because the GEI variables from Sim1 did not account for directional relationships, as GEI variables were independently sampled from the general covariance matrix. In contrast, Sim2 positioned the environments according to their similarity in the climate context of that given generation. This difference can be quantified in the Euclidean distance relationship for the entire generation. Table 6 shows that the mean distances correctly indicated that E1 and E2, and E3 and E4 were closely related to each other. The SD were below 0.06 in the Sim1 but ranged from 0.18 and 0.40 in Sim2, reflecting generation-specific variation in environmental variables. Since environmental variables were generated (sampled) again each generation, Euclidean distances between environments were expected to vary accordingly. In this sense, the larger SD in Sim2 indicate that it regenerated GEI, accounting for generation-specific environmental variation. Despite intergenerational deviations, the overall relationships among environments were captured by the environmental covariance structure, with the mean Euclidean distance effectively summarizing these relationships (Table 6). The Sim1 was inappropriate for GEI graphical interpretation, as it did not reveal a ‘which-won-where’ pattern (Seyoum 2021). This is critical because this limitation can distort selection decisions and lead to inaccurate ASV calculations, which determine the genotype stability across the environments. On the other hand, Sim2 results aligned with other findings in visualization studies utilizing biplots, which showed closer distances between environments with similar conditions and captured directional relationships (Vaezi et al. 2017; Yan et al. 2007). These directional relationships are critical for genotype recommendations in specific mega-environment (Yan and Tinker 2005).

**Table 6.** The Euclidean distance between a pair of environments under two simulation approaches (Sim1 and Sim2) in biplot. Values represent mean ± SD across 50 replicates.

| Env<sup>†</sup> \ GEI var | Sim1 |  |  |  | Sim2 |  |  |  |
| --- | --- | --- | --- | --- | --- | --- | --- | --- |
|  | 0.1 | 0.5 | 1 | 2 | 0.1 | 0.5 | 1 | 2 |
| E1-E2 | 1.04±0.01 | 1.04±0.02 | 1.04±0.02 | 1.04±0.01 | 0.94±0.38 | 0.96±0.41 | 0.94±0.40 | 0.96±0.39 |
| E1-E4 | 0.97±0.01 | 0.97±0.01 | 0.97±0.01 | 0.97±0.01 | 1.17±0.21 | 1.16±0.22 | 1.16±0.21 | 1.18±0.20 |
| E1-E3 | 1.41±0.00 | 1.41±0.00 | 1.41±0.00 | 1.41±0.00 | 1.22±0.21 | 1.25±0.19 | 1.25±0.19 | 1.23±0.20 |
| E2-E4 | 1.41±0.00 | 1.41±0.00 | 1.41±0.00 | 1.41±0.00 | 1.18±0.24 | 1.16±0.24 | 1.18±0.25 | 1.16±0.26 |
| E2-E3 | 1.03±0.01 | 1.03±0.01 | 1.03±0.01 | 1.03±0.01 | 1.20±0.19 | 1.19±0.18 | 1.18±0.20 | 1.18±0.21 |
| E3-E4 | 0.96±0.02 | 0.96±0.02 | 0.96±0.02 | 0.96±0.02 | 0.99±0.38 | 0.95±0.41 | 0.98±0.39 | 0.97±0.38 |

<sup>†</sup>Pairs of assessed environments (e.g., E1-E2, E1-E3, etc.)

To further support this idea, correlations of ASV across all generations were compared between Sim1 and Sim2. In this study, lower correlations were observed with decreasing GEI variance, with correlations of 0.77, 0.70, 0.68, and 0.67 corresponding to GEI variances of 0.1, 0.5, 1, and 2, respectively. These results indicate that the Sim1, which did not account for directional relationship, distorted the stability trend. In other words, the application of Bayesian AMMI effectively provides graphical representations that reflect the genotype stability by integrating the generated TGVs and environmental variables.

### Utilization of simulation framework

Applying the Bayesian AMMI framework provides two novel insights for GEI simulation: (1) Integration of high-dimensional environmental covariates. The computational cost of building **Ω** is not strongly dependent on the number of environmental covariates. In this sense, the more complex high-dimensional environmental covariates can be considered for GEI simulation study. This allows researchers to account for realized interactions among multiple environmental conditions, enabling the generation of more complex and realistic environmental scenarios. (2) Simulation of directional relationships between genotypes and environments. The Bayesian AMMI approach captures directional relationships accounting for GEI by incorporating pre-simulated variables specific to the given generation. This facilitates the investigation of genotype stability and its variation under different breeding strategies. While this simulation framework enables the generation of GEI variables with appropriate directional relationships, future studies should extend the framework to account for multiple chromosomes.

In conclusion, the Bayesian AMMI framework, incorporating high-dimensional environmental covariates, effectively captured directional genotype-environment relationships, producing biplot representations that accurately reflect environmental similarity. This framework enables appropriate simulation of GEI accounting for complex environmental conditions. Based on these findings, we propose using a Bayesian AMMI framework for GEI simulation, with future development needed to extend its application to multiple chromosomes.

## Data availability statement

The pipeline for simulating described genotype-by-environment interactions is included in Supplementary File 1 to Supplementary File 3. The authors affirm that all data necessary for confirming the conclusions of the article are present within the article and figures.

## Acknowledgements

The authors thank members of the Jarquin and Hidalgo research groups for helpful discussions and feedback during the development of this study. This research did not receive any specific grant from funding agencies in the public, commercial, or not-for-profit sectors.

